# An interactive viral genome evolution network analysis system enabling rapid large-scale molecular tracing of SARS-CoV-2

**DOI:** 10.1101/2020.12.09.417121

**Authors:** Yunchao Ling, Ruifang Cao, Jiaqiang Qian, Jiefu Li, Haokui Zhou, Liyun Yuan, Zhen Wang, Guangyong Zheng, Guoping Zhao, Yixue Li, Zefeng Wang, Guoqing Zhang

## Abstract

Comprehensive analyses of viral genomes can provide a global picture on SARS-CoV-2 transmission and help to predict the oncoming trends of pandemic. This molecular tracing is mainly conducted through extensive phylogenetic network analyses. However, the rapid accumulation of SARS-CoV-2 genomes presents an unprecedented data size and complexity that has exceeded the capacity of existing methods in constructing evolution network through virus genotyping. Here we report a Viral genome Evolution Network Analysis System (VENAS), which uses Hamming distances adjusted by the minor allele frequency to construct viral genome evolution network. The resulting network was topologically clustered and divided using community detection algorithm, and potential evolution paths were further inferred with a network disassortativity trimming algorithm. We also employed parallel computing technology to achieve rapid processing and interactive visualization of >10,000 viral genomes, enabling accurate detection and subtyping of the viral mutations through different stages of Covid-19 pandemic. In particular, several core viral mutations can be independently identified and linked to early transmission events in Covid-19 pandemic. As a general platform for comprehensive viral genome analysis, VENAS serves as a useful computational tool in the current and future pandemics.

## Background

The World Health Organization (WHO) characterized Coronavirus disease 2019 (COVID-19) as a global pandemic (pandemic) on March 11, 2020. By the end of October, more than 45 million COVID-19 cases had been diagnosed with > 1 million deaths (https://covid19.who.int/). More than 190,000 SARS-CoV-2 genomes have been sequenced and published (https://www.epicov.org/epi3/) since the end of 2019. The comprehensive analyses of these genomes could provide a global picture of how the virus transmitted among different populations, which may help predict the oncoming trends of pandemic. The main approach for molecular tracing of viral transmission is to thoroughly compare the genomes of different viral strains, leading to a series of phylogenetic trees or evolution networks that can also help to interpret the genomic mutations along transmission [1–3].

Previously Lu and colleagues used linkage disequilibrium and haplotype map to analyze the genomes of 103 SARS-CoV-2 samples, and classified the viral genomes into type L and S based on mutations at positions 8782 and 28144 [4]. In another report, Corlett et al. used phylogenomic analyses to classify 93 SARS-CoV-2 genomes into 58 haplotypes, and further clustered these virus strains into five clades [5]. Similarly phylogenetic network was also constructed by Forster et al. using 160 viral genomes, which classified the viruses into three types (A/B/C) based on the nucleotide variants at loci 8782, 26144, 28144, and 29095 and the corresponding changes of viral proteins [6]. In these evolutionary analyses, the R package Pegas and the GUI-based software DNASP and POPART were used due to their user-friendly interfaces and the convenience of programming [7–9]. However, construction and interpretation of the viral genome evolution network had become increasingly complicated with rapid accumulation of available viral genomes, which make it difficult to artificially genotype and cluster the viruses from the network. In addition, since the construction of evolution network involves extensive calculations of large matrices, single-threaded analysis tools are unable to complete the analysis within a reasonable time limit in order to continuously track the dynamic changes of viral mutational patterns.

To meet these challenges, we developed a new software pipeline, the viral genome evolutionary analysis system (VENAS), to integrate the analyses of genomic variations and evolution networks. We applied minor allele frequencies (MAFs) of the effective parsimonious information sites (ePISs) on viral genomes as weighs to calculate the Hamming distance matrix between different genome sequences [10, 11], and used the neighbor-joining method to build the evolution network. We adopted highly paralleled multi-processing strategy to maximize the computation efficiency in calculating the distance matrices. The VENAS further applied a community detection method to transform the evolution network into a two-dimensional isomorphic topological space, with a unique PIS pattern in each sub-space. Finally, we used the network disassortativity trimming algorithm to extract the backbone network of the topological space, which can be used to trace the evolution of viruses and detect core mutations among distinct strains. The development of VENAS enables researchers to rapidly construct large-scale evolution network at a reduced time and computational cost, and explore the topological space in an interactive interface.

## Results

### Design principle and construction of VENAS

The VENAS seeks to apply reliable computational algorithms to build an integrative genomic analysis system that enables researchers to trace viral mutations along the transmission routes using the daily updated SARS-CoV-2 genomes. The construction of VENAS is illustrated in figure 1. Briefly, we first collected all available SARS-CoV-2 genome dataset with stringent quality-control (Fig 1a). Each genome was assigned with a series of binary labels (PIS, ν) that identify the parsimonious information sites (PISs) and their corresponding allele frequencies, where a unique set of PISs present a distinct genome type. The PISs with frequencies above an user-defined threshold (i.e., effective PISs, or ePIS) were used to calculate the Hamming distance between distinct genome types [12] (Fig. 1b).

**Figure 1:**
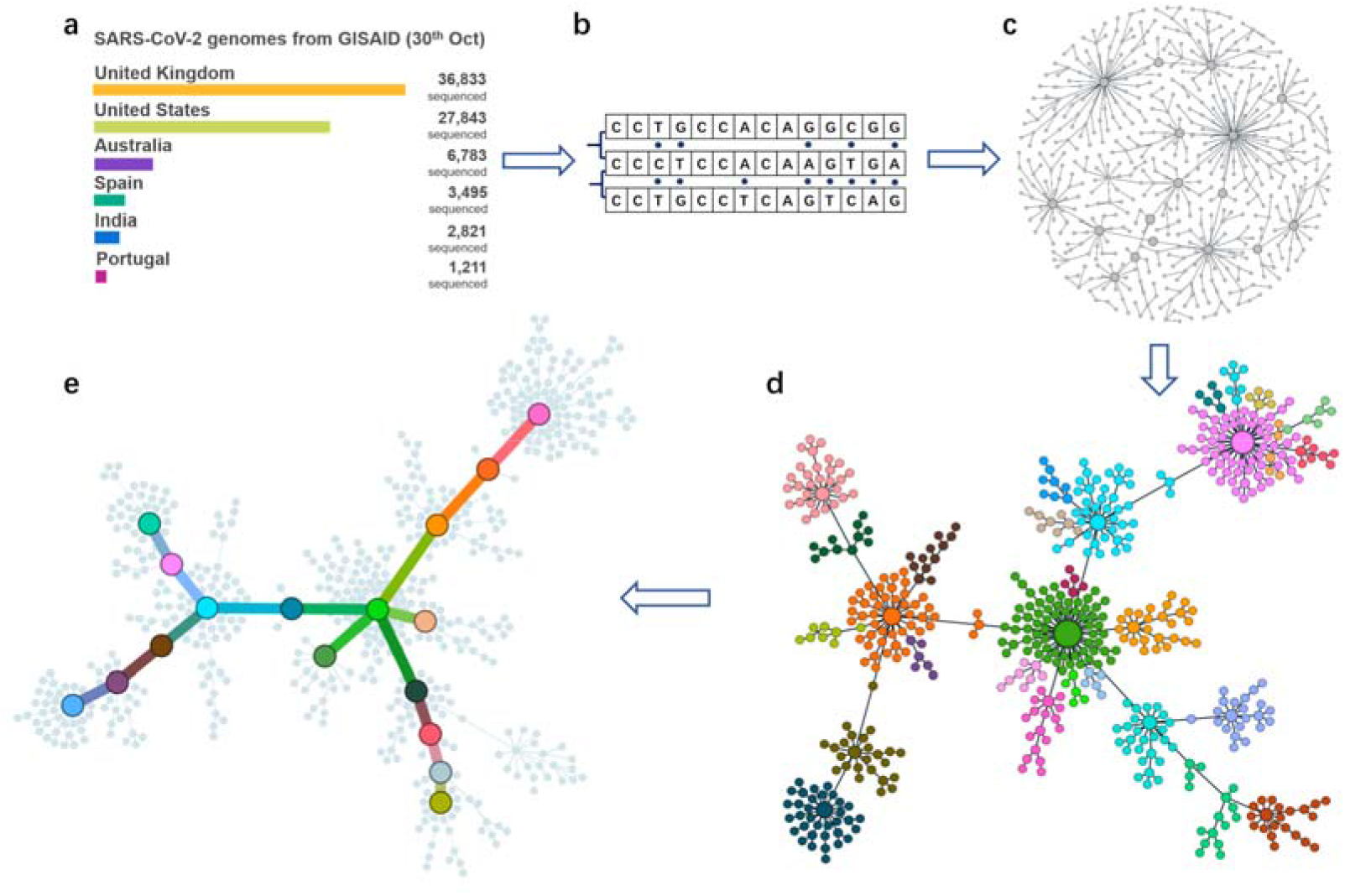
The construction of VENAS. a) Collecting a large number of SARS-CoV-2 genomes from the GISAID database, the genomes with same set of PIS were assigned as the same types; b) Generating a distance matrix between different genome types based on the Hamming distance, sorting and adjusting the distances with MAFs; c) Constructing the viral evolution network for genomic variation analysis; d) Topologically clustering the network into subspaces using community detection; e) Analyzing the evolution network to extract the major transmission paths that consist of core genome types with associated core mutations.

We further sorted the distances between different genome types in an ascending order and sequentially connected all types using the neighbor-joining method, resulting a fully connected viral genome evolution network with the shortest sum of distance (Fig. 1c, the nodes present genome types and the edges present adjusted Hamming distances). During this process, the pairs of genome types with the same distance were adjusted by the MAFs of differential ePISs, where the pairs with lower MAFs (i.e. higher conservation) were given higher rank (see methods). With the assumption that the number of genomic mutations occurred during a single human to human transmission is small, the neighbor-joining method can effectively combine viral genomic alterations with transmission events to reliably construct a network of viral evolution. Unlike the phylogenetic tree, we did not limit with bifurcation connection, thus giving additional spatial freedom to the network that is more consistent with the multi-dimensional transmission patterns of the virus (Fig. S1).

We next used a disjoint community detection method to cluster the evolution network into topologically linked subdomains, which represent different evolution clades containing many closely-connected genome types (Fig. 1d). Such segmentation enable us to subjectively identify the topological clades with “tight” intraclade connectivity and the “sparse” interclade connectivity, which reflect the relationship of different genome types among viral communities formed through the natural transmission. Finally, we used the network disassortativity trimming algorithm to extract the core genome types (i.e., nodes) from the evolution network, and further calculated the shortest paths between the core nodes using the Dijkstra algorithm [13], generating a “backbone network” that recapitulates the main mutational paths (Fig. 1e). Many SARS-CoV-2 genome samples also contain epidemic information associated with the cognate patients, which was further integrated into the VENAS network. Integration of epidemic information with the virus genome type allowed us to directly infer the viral evolution patterns at critical points of transmission. With the large sample numbers and the robust community detection algorithm, the core nodes in VENAS network can be identified despite some uneven sampling or missing samples.

The users can visualize the viral genome evolution network of VENAS using a general relationship graph or force-directed graph tools, such as the web-based Apache Echarts (https://echarts.apache.org/), d3.js (https://d3js.org/), or the application-based Gephi.

### Efficient network construction with parallel computing

The computation of MAF-adjusted Hamming distance matrices is the rate limiting step during the construction of viral evolution network of VENAS. To improve the computing efficiency, we took advantage of the multi-core and multi-threaded features in high-performance computers and developed a highly parallel network construction pipeline (see methods). To compare the computational efficiency of VENAS with previous tools, we used three SARS-CoV-2 genomic datasets with sample numbers at different orders of magnitude (103, 1,050 and 14,949 samples respectively), and analyzed them using VENAS, Pegas, and POPART, respectively. These pipelines were tested using the same hardware environment (Dual Intel^®^ Xeon^®^ Gold 6129 processors with 32 cores and 64 threads, 768GB memory) and similar program compilers (PyPy 7.3.0 with GCC 7.3.1 for VENAS v1.0; R 3.6.1 for Pegas v0.13; and Qt 4.8.4 for POPART v1.7). For each software, we evaluated their capacity to handle large datasets and their computation time (Table 1).

**Table 1.** Virus evolution network benchmark of Pegas, POPART and VENAS

| Dataset | Viral genome<br>samples | Nodes | Links | Cal. Duration |  |  |
| --- | --- | --- | --- | --- | --- | --- |
|  |  |  |  | Pegas | POPART | VENAS |
| Dataset 1 | 103 | 34 | 33 | 28s | 1.45s | 0.48s |
| Dataset 2 | 1,050 | 300 | 299 | 56s | 5min | 3s |
| Dataset 3 | 14,949 | 7,846 | 7,845 | NA(>48h) | NA(>48h) | 10min |

Note: Since PEGAS and POPART do not include the topological clustering and backbone network extracting functions, the computing efficiency comparison between the three software/toolkits only includes the common Hamming distance matrix operations and the network construction processes.

We found that all the three software could efficiently compute the evolution network containing hundreds of samples (dataset 1), with VANAS finished the task in 0.48 sec (near real-time computation) whereas the Pegas takes ~28 sec. With the increasing of the dataset scale into thousands of samples, all the three software could still complete the computational task within a reasonable amount of time. The VENAS achieved best performance by finishing the computation in 3 sec, with 18x and 100x acceleration compared to POPART and Pegas, respectively. When the number of genomes reached to tens of thousands (dataset 3), neither Pegas nor POPART could complete the computation in a finite time (no detectable progress in 48 hours), probably due to the single-threaded computing limitation. However, VENAS fully used all the 64 threads in the hardware environment completed the computation in ~10 minutes, making it possible to handle data accumulation in the future

### Comparison of the VENAS networks with previous results

As mentioned earlier, evolution networks have an advantage over phylogenetic trees in terms of the spatial freedom of connectivity. Comparing with previous software (e.g., POPART and Pegas) that construct phylogenetic network using haplotypes, VENAS generates evolution network based on the MAF-adjusted Hamming distances between different genome types, allowing automatic identification of major transmission paths and core genome types without manual clustering. As a validation, we first examined if the VENAS can recapitulate the SARS-CoV-2 haplotype network identified by an early study using a small number of genomes before March 2020 [4]. Two clusters of SARS-CoV-2 bearing core variations in loci 8782 and 28144 were previously identified as the types L (8782C/28144T) and S (8782T/28144C), and the community detection method in VENAS can reliably separate the L/S type into two clades with the same core mutations (comparing left and right panels in Fig. 2a).

**Figure 2.**
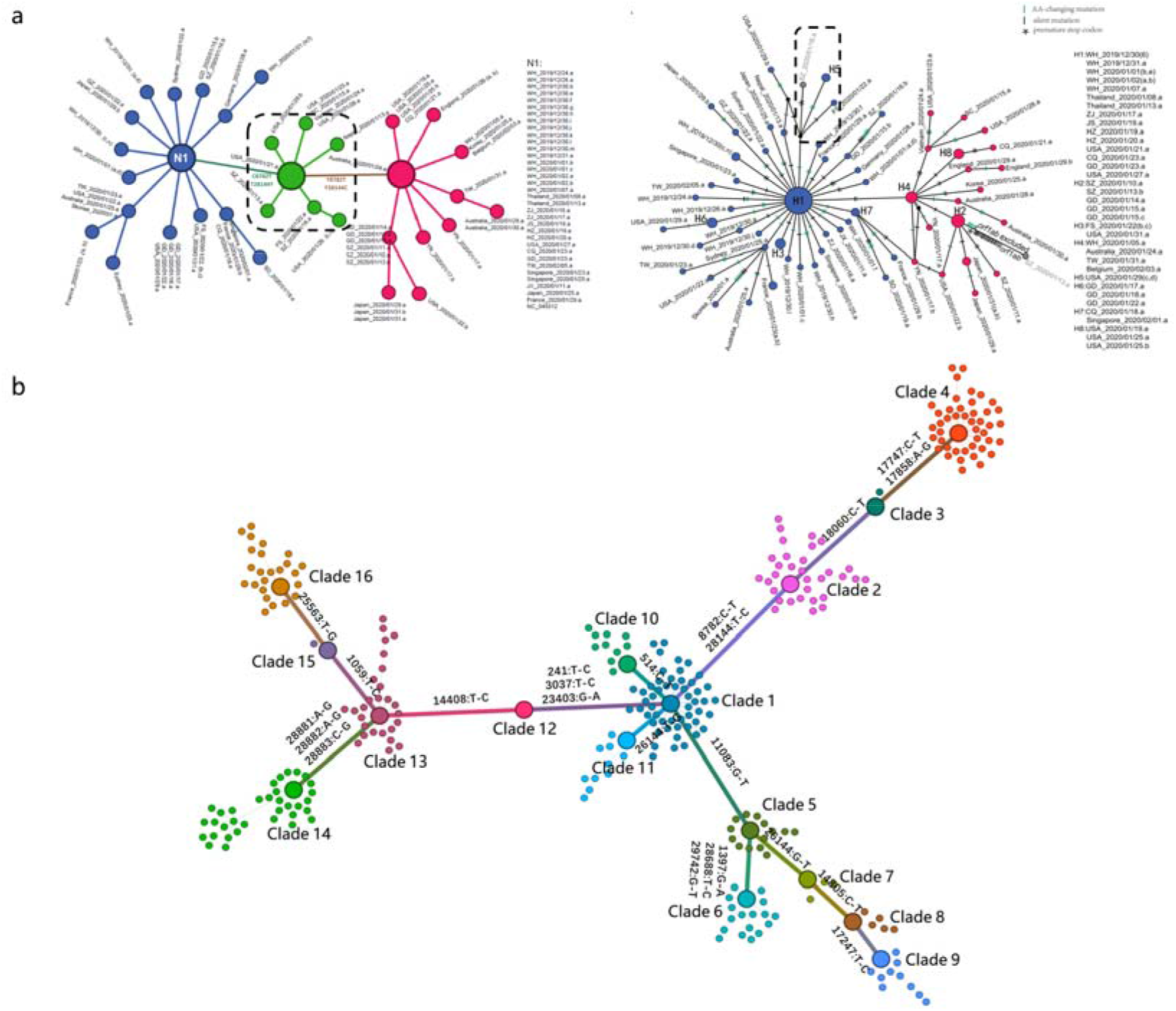
VENAS recapitulate the key transmission route identified from an earlier study. a) VENAS recapitulate the key transmission route identified from an earlier study. Left, regenerating the evolution network of early transmission using VENAS pipeline and the dataset in Lu’s article containing 103 viral genomes and 34 genome types (dataset 1). Right, the original haplotype network plotted in Lu’s article for comparison; b) VENAS evolution network generated from SARS-CoV-2 genomes publicly available before March 25, 2020 (dataset 2), containing 1050 viral genomes and 546 genome types; the network is clustered into 16 topological clades, with self-increasing ordinal numbered clade names, and labelled links with differential variations between clades.

More importantly, VENAS also identified an intermediate clade of the two types (top panel of Fig. 2a). This new clade contains the core variant 8782Y/28144Y with T or C at each position, which could be potentially caused by a co-infection of both types in the corresponding patient (USA 2020/01/ 21.a, GISAID ID: EPI_ISL_404253) who visited Wuhan in December 2019 and returned to Chicago in January 13 2020. Alternatively, the transition between the type L and S may have occurred during the infection of this patient, who would then be a founder of the new viral type. In brief, by combining the core genomes types and corresponding variations from the massive amount of viral genomes, VENAS enables researchers to explore the major transmission paths with improved resolution and an informative graph.

### Topology-based detection of core variations at early stage of global transmission

We further expanded the VENAS to analyze the viral genomes at the beginning of global transmission using a dataset containing all entries publicly released by March 25, 2020. The evolution network containing 1,050 viral genomes were constructed by VENAS in three seconds, producing a “backbone network” with 16 major viral clades in the topological space that reflect the possible transmission paths (Fig. 2b). Each pair of viral clades was separated by a group of core variants that also were identified by comparing PIS sites in these clades (Table 2). Some clades and associated variants were also reported by other researchers independently. For example, the core variants C8782T/T28144C recognized as the PIS between clades 1 and 2 (i.e., separating the clades 2, 3 and 4 from the other clades), which was also reported to distinguish the “type L/S” [4, 6]; The variants G11083T between clades 1 and 5 were first identified from the patients in Diamond Princess Cruise and were also reported by Sekizuka et al [14]. Some of the clades also provided new information on virus transmission. For example, the clades 1 and 12 were connected by a set of tightly-linked variants (C241T/C3037T/A23403G), which can cause single amino acid substitution in S protein. This mutational event coincided with the early transmission to Europe as it was first identified in a patient traveled from Shanghai to Bavaria for a business meeting of Webasto[15].

**Table 2.** Summary of core variants at the early stage of global transmission.

| No. | Position/ | Class (protein changes) | First identified date | Ref. |
| --- | --- | --- | --- | --- |
| 1 | T28144C/C8782T | ORF8:p.L84S/nsp4:p.S76S | Jan. 05, 2020<br>Wuhan/WH04 | [4, 6] |
| 2 | C18060T | nsp14:p.L7L | Jan. 19, 2020<br>USA/WA1 | [16] |
| 3 | C17747T/A17858G | nsp13:p.P504L/nsp13:p.Y541C | Feb. 20, 2020<br>USA/WA-S2 | [17] |
| 4 | G11083T | nsp6:p.L37F | Jan. 18, 2020<br>Chongqing/IVDC-CQ-00<br>1 | [14] |
| 5 | G26144T | ORF3a:p.G753V | Jan. 25, 2020<br>Hangzhou/ZJU-01 | [6] |
| 6 | C14085T | nsp12:p.N215N | Feb. 27, 2020<br>USA/WA3-UW1 |  |
| 7 | T17247C | nsp13:p.R337R | Feb. 28, 2020<br>Brazil/SPBR-02 |  |
| 8 | G1397A/T28688C/G29742T | nsp2:p.V378I/N:p.L139L | Jan. 18, 2020<br>Wuhan/HBCDC-HB-05 |  |
| 9 | T514C | nsp2:p.H83H | Mar. 02, 2020<br>Netherlands/Hardinxveld<br>_Giessendam_ 1364806 |  |
| 10 | A23403G/T241C/T3037C | S:p.D614G/ | Jan. 28, 2020<br>Germany/BavPat1 | [18, 19] |
| 11 | C14408T | nsp12:p.P323L | Feb. 20, 2020<br>Italy/CDG1 | [18] |
| 12 | G25563T | ORF3a:p.Q57H | Feb. 26, 2020<br>France/PL1643 | [20] |
| 13 | C1059T | nsp2:p.T85I | Mar. 09, 2020<br>France/PL1643 |  |
| 14 | G28881A/G28882A/G28883C | N:p.R203K/N:p.G204R | Feb. 25 2020<br>Germany/Baden-Wuertt<br>emberg-1 | [21] |

Identification of the previously reported key mutations/variants suggests that VENAS can serve as a reliable analysis pipeline for the research community. In addition, VENAS also identify novel mutations that function as key evolution markers to separate the viral clades. For example, when expanding VENAS to analyze a large-scale dataset of 56,768 SARS-CoV-2 entries in the end of October 2020, we identified several additional variations that separate key clades, including variants that changed amino acid sequence of important viral proteins (such as C22227T and C21614T in S protein, C28932T in N protein, G29645T in ORF10). Further study of these variants may reveal the evolution and transmission patterns of the virus in the late stage of the epidemic. By linking the molecular level variations (i.e., changes in nucleotides and amino acids) with the epidemic information (e.g., patient population and their travel and contact history), the researchers may discover new clues for viral evolution and transmission.

### Tracing the viral evolution network through different stages of transmission

To compare the mutational features of viral genome during different periods of COVID-19 pandemics, we applied VENAS to analyze the cases collected in different times, resulting viral evolution networks at different pandemic stages (Fig. 3). We found that, during the early stage of global pandemic (before April, 2020), the viral genome types showed a limited diversity and can be clustered into 16 main clades (Fig. 3a). These clades were centered around a clade containing the earliest strain detected mainly in Wuhan (clade 1). Importantly, the branching pattern of this evolution network reflected a clear path of transmission. For example, the “North American branch” (clade 2 -> 3 -> 4) containing variation C17747T/A17858G was directed branded out from clade 2, which was also named as type S previously[4, 17]. The viruses in the “Diamond Princess branch” (clade 5 to 9), carried by most patients in the community transmission event of the cruise ship, contain a key variation (G11083T) that caused an amino acid substitution in NSP6 protein. Finally, the “European branch” containing variation A23403G/C14408T was further divided into two branches characterized by variations G25563T and G28881A/G28882A/G28883C (clade 14), which were wide spread in Europe and North/South America.

**Figure 3.**
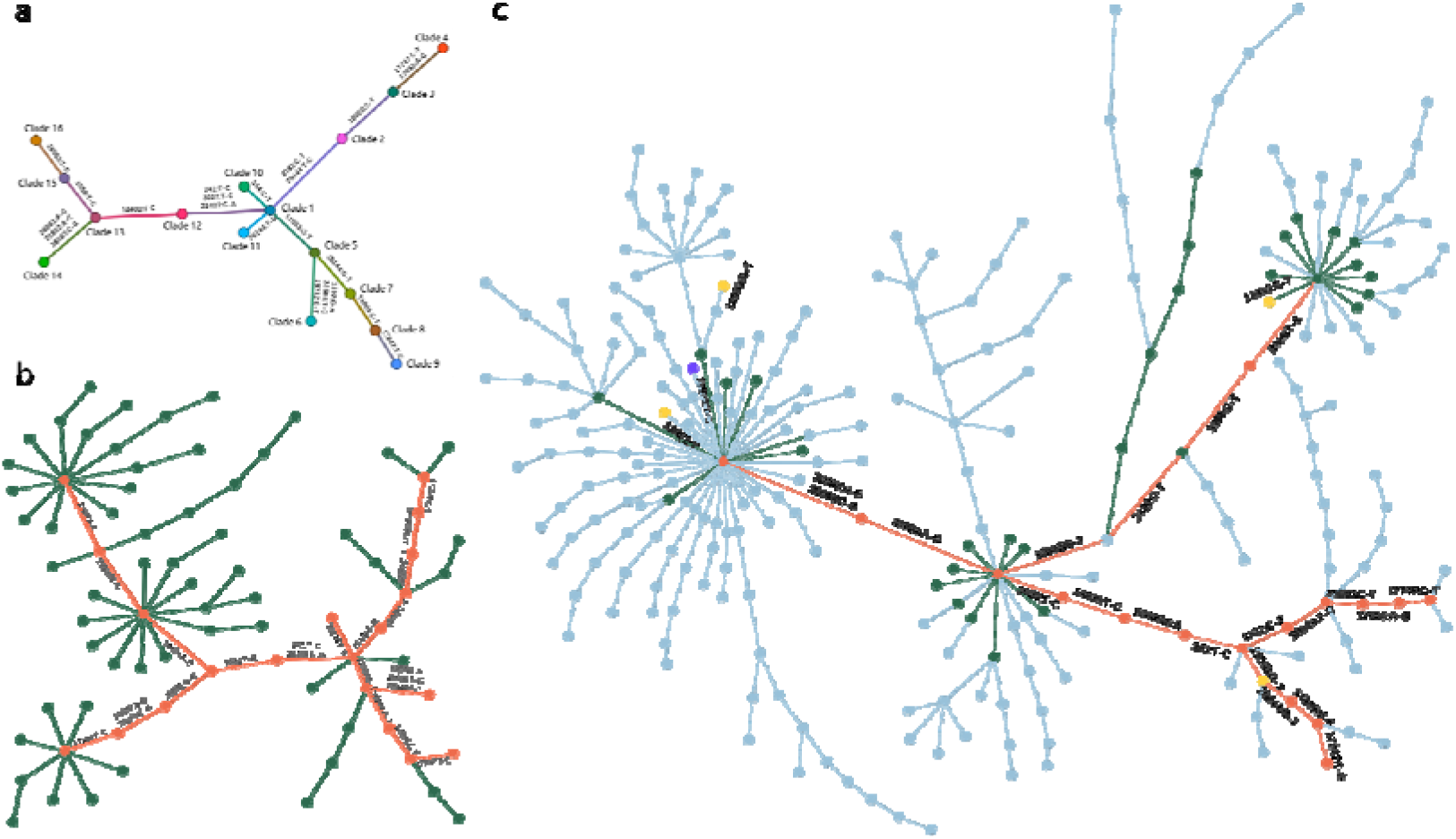
VENAS networks at different SARS-CoV-2 pandemic stages. a) The VENAS network with key mutations using the genome data by 3/25/2020. The 16 clades are numbered as Fig. 2b. b) The VENAS network with key mutations using the genome data by 6/30/2020. The earlier clades identified were labeled in color orange, and the new clades were labeled in green. c) The VENAS network with selected key mutations and recurrent mutations using the genome data by 10/30/2020. The old clades were labeled with same colors (orange and green), while new clades developed since June were labeled in light blue.

In the mid-stage of the pandemic (pre-June, 2020), the viral genome types expanded into additional branches with increasing diversity, and the cases in some of the original branches showed a slower growth compared to the new branches (Fig. 3b and Supplementary table S3). With the global transmission of the SARS-CoV-2, we found that the original “European branch” mutated into additional genome variants, resulting in many novel types that were clustered into new clades and sub-branches. As a comparison, the original “Wuhan clade” branched slowly with a much smaller genome type diversity, probably due to successful confinement of the virus in China. Meanwhile, some of the clades showed a linear sequential branching pattern with small number of variations detected (e.g., the Princess Diamond cruise branch), likely reflecting a good prevention and control of virus transmission.

By the late stages of the pandemic (before October, 2020), the network continued to branch out into new clades containing additional genome types, whereas the original clades become less prominent (Fig. 3C and Supplementary table S4). However, it became difficult to define a clear population profile and transmission paths path for each clade. The cases from Europe and North America population are dominant in almost every clade, probably due to the large patient numbers and the extensive sequencing. In addition, several variations (e.g. G11083T, C21757T) were detected in multiple viral genome types from different clades, reflecting recurrent mutations independently evolved from multiple places without direct transmission. These genomic regions may be the mutational “hot-spots” that should be closely monitored in the further.

## Discussion

To meet the challenges of the massive amount of sequencing data accumulated during COVID-19 pandemic, we developed a novel method for genomic tracing of virus transmission through an integrative analysis of viral genome and epidemic information. Several new algorithms were implemented in a high-performance parallel computing environment, resulting a rapid construction of high-resolution evolution networks of SARS-Cov-2 with interactive interface. Previously, the common method to study viral genome variations is through construction of phylogenetic trees, often using a neighbor joining algorithm [22]. We made important improvement in VENAS by changing the binary structure of the phylogenetic trees into a multi-dimensional force-directed graph, and adjusting the order of ambiguous connections with MAFs. The VENAS effectively clustered the viral genome types into clades by the topological layout of the graph, which reflected the major evolutionary patterns between virus genomes within each clade and provided a basic data model for the study of virus transmission trends.

To reduce the artifact from sequencing errors, we used PISs instead of all variation sites to compute the evolution network. The PISs were further filtered by a threshold of variant frequency to obtain ePISs, which enhanced the confidence level and the direct comparison of MAFs even with incomplete sequencing. Since all the SARS-CoV-2 genomes are extremely similar, thousands of pairs of genome types have the same Hamming distance, and thus the order of connections may greatly affected the reliability and precision of the network. VENAS combined a biological measurement of variation frequency (MAF) with the Hamming distance by assigning the MAF as a weight of Hamming distance, and thus the differential genome types with the maximum conservation were firstly connected to enhance the robustness of the network. In addition, sequencing of viral samples often produces ambiguous bases in differential PISs (i.e. degenerate bases like Y, R, or N). While the Pegas skips the links with ambiguous bases and the POPART directly deletes all relevant viral genome, the VENAS make optimizations in data quality control and subsequent algorithm. The sequences containing ambiguous bases in high-MAF ePISs were selectively and optionally removed to ensure the precision of the major paths of the network.

It should also be noted that the VENAS network is a non-directional acyclic graph, and thus people should combine the epidemiological information (like time of sampling) when inferring transmission path from the network. Another caveat during network interpretation is that the “variant reversion” could be mistakenly called in some areas because the neighbor-joining method tend to force a fully connected network despite missing samples. In such case, a base at the certain locus may appear to be mutated back and forth on two adjacent or nearby edges. The insufficient sampling of some specific genome types can lead to a missing node that is required to construct a coherent path of nodes reflecting the real transmission events. In such situation, the algorithm will “enforce” the network construction through a neighbor node that is the closest to the real node.

The enormous amount of mutational data accumulated during viral transmission was manifested in VENAS as large community-based clades through the modularity-based community detection algorithm such as Louvain. In each topological clade identified, the central node represents the predominant genome type in the given viral community, and the genome diversity of the viral community was reflected by the numbers of the inward and outward edges. We employed the network disassortativity trimming algorithm to merge the small clades at the edge of the network with the major clades, and the resulting backbone network of the major clades can effectively reflect the major evolutionary paths and associated core variations.

In conclusion, the genomic tracking is critical to understand the global transmission of the SARS-CoV-2. We developed a software platform that can handle the massive amount of viral genomic data and rapidly generate the evolution network using a highly paralleled computation protocol. The topology-based community detection and the network disassortativity trimming algorithm of this method also enabled the identification of critical viral groups that formed a backbone network of virus transmission. Using the interactive user interface of VENAS, we were able to identify several known branches in the early stage of COVID-19 pandemic. We also detected additional new branches and sub-branches that help to interpret the transmission trends of later stages. As a general platform, we expect that VENAS can serve as a valuable tool for data analysis and visualization that support virus tracing and drug development for further pandemics.

## Methods

### SARS-Cov-2 genome datasets

Full-length viral genome sequences were collected for genomic comparison and network construction. In this study, we selected four datasets from different stages of the COVID-19 pandemic. The first dataset cited 103 SARS-CoV-2 genome sequences used in the research by Lu et al. [4](Supplementary table S1). The second dataset selected viral genomes published in the early stage of the pandemic by March 25, 2020, with 1,050 left after quality assurance steps (Supplementary table S2). The third dataset contained 14,949 SARS-CoV-2 genomes sequences released in the mid stage of the outbreak by June 30, 2020 (Supplementary table S3). The fourth dataset included all 56,768 high-quality viral genome sequences by October 30, 2020 (Supplementary table S4). All datasets are derived from publicly available SARS-CoV-2 (taxon_id=2697049) genome sequence data from GISAID and Genbank.

### Effective parsimony-informative site

The calculation of Hamming distance matrices was based on the number of differences in the effective Parsimony-Informative Sites (ePISs). First, all SARS-CoV-2 sequences should be multiple-aligned using MAFFT [23] to obtain a consensus alignment in ma format. Next, the number of unambiguous bases for each variation loci in all genomes was summed up, filtered and finally obtained the ePIS list. The filtering strategy was as following: 1) The site was a parsimony-informative which contained at least two types of nucleotides, and at least two of them occur with a minimum frequency of two; 2) the PIS was effective if the number of unambiguous bases ≥80% of the total genomes.

### Minor allele frequency

MAF (Minor Allele Frequency, ν) a quantitative indicator to evaluate the rareness of variations. We calculated each ePIS and used the maximum MAF of differential ePISs between two viral genomes to adjust the Hamming distance. Since ambiguous bases such as N, -, and degenerate bases increased the uncertainty in the later computation of Hamming distances, VENAS users could use MAF to filter viral genomes with ambiguous bases in high MAF ePISs.

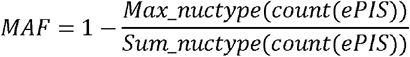

### Viral genome evolution network construction

Firstly, we gave a set of ePISs to each viral genome and used this combination of ePIS (genome type) to represent it. Therefore, the number of genomes was reduced by eliminating redundancy genome types. The MAF(ν) of each ePIS was also calculated and indexed for later queries. Then we calculated the Hamming distance (d) matrix [12], which defined the viral phylodynamic relationship between each two genome types[24, 25], to evaluate the evolutionary distances. The Hamming distance matrix M(d_ij_, νmax_ij_) was formed by combining the distance values with the maximum MAF of differential ePISs (Fig. S2a). If the pair of genome types contained ambiguous bases, the distances of those ePISs were calculated as 0.

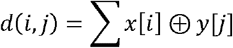

Secondly, we constructed the evolution network by using the MAFs adjusted Hamming distance matrix. All pairs of genome types in M(d_ij_, νmax_ij_) were sorted by d_ij_ and νmax_ij_ in ascending order, resulting in a sorted Hamming distance list L(d_ij_, νmax_ij_) (Fig. S2b). Then pairs of genome types were connected in sequence according to the list (Fig. S2c). At each step of the connection, we abandon the link if the two genome types (P_i_ and P_j_) associated with the d_ij_ were already indirectly connected by other genome types (Fig. S2d). Finally, the fully-connected evolution network G(P,d) was constructed, with the nodes representing the genome types and the edges representing the differential ePISs (Fig. S2e).

### Topological classification and major path recognition

Two third-party Python libraries, networkx (https://networkx.github.io/) and CDlib420(https://github.com/GiulioRossetti/CDlib), were used in the topological clustering and backbone network extracting process once the viral genome evolution network was constructed.

Firstly, we used the Louvain modularity community detection algorithm to cluster the evolution network to a set of topological clades[26]. The Louvain algorithm [27] calculated the gain in modularity [28] as an optimization target and used a heuristic method to find the community clusters. Compared to other clustering algorithms, Louvain was faster to compute on large graph networks with guaranteed accuracy.

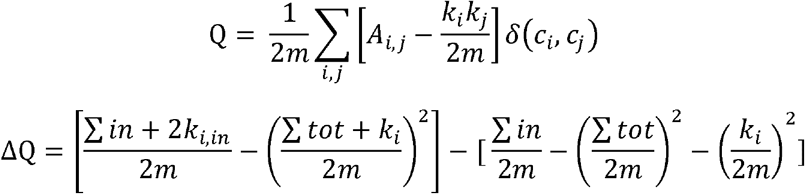

Secondly, we applied a network disassortativity trimming algorithm to extract the core viral genome types of the evolution network. A highly diverse viral genome type tended to have a higher inward/outward degree as the edges represented the possible evolutionary paths of the virus. We analyzed the statistical degree distribution of all nodes from the network. The vast majority of the nodes had a degree of 1 (Fig. S3). Therefore, we removed the nodes with degree 1 from the network and then filtered the remaining candidate nodes using the network disassortativity trimming algorithm [29] to obtain the core viral genome types.

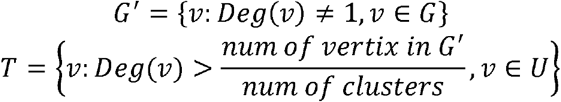

Finally, all core viral genome types were selected to complete the identification of the backbone network. We employed the Dijkstra algorithm to find the shortest path between each pair of genome types. All nodes on the shortest paths, no matter the core node or not, were added to the list of core viral genome. Thus the complete evolutionary paths between the core viral genome types and corresponding core variations were discovered. We integrate all the major evolutionary paths to complete the identification and construction of the backbone network.

## Ethics approval and consent to participate

Not applicable.

## Consent for publication

Not applicable.

## Availability of data and materials

VENAS is cross-platform open-source software available at https://github.com/qianjiaqiang/VENAS. It can be easily installed from source code or using pre-build binary.

## Acknowledgments

The author thanks the GISAID EpiFlu™ Database, the NCBI Genbank Database, and all contributors for sharing SARS-CoV-2 genome sequences and corresponding metadata.

---

**Figure S1.**
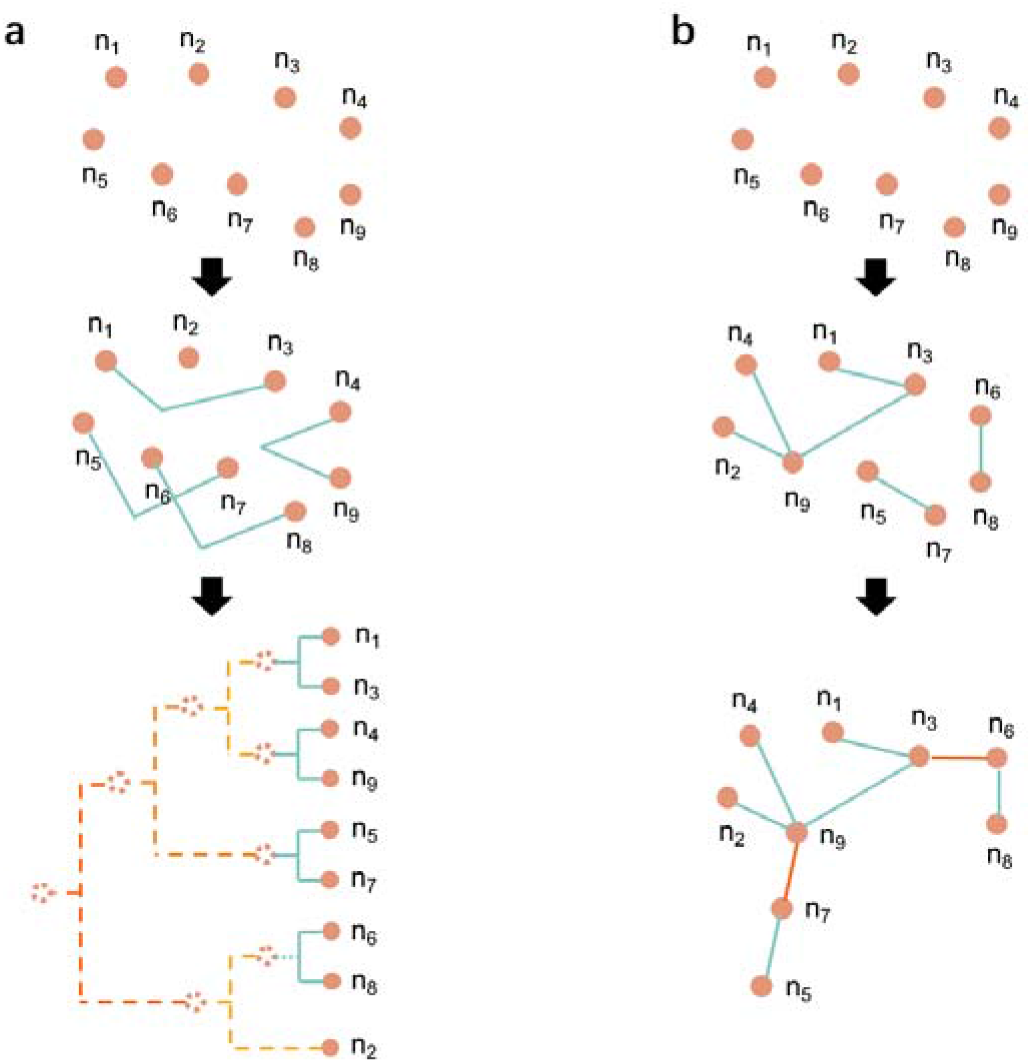
The comparison between the construction processes of the phylogenetic tree and evolution network. a) The construction process of the phylogenetic tree using the neighbor-joining method. Virtual nodes representing the consensus of child nodes were generated to connect the binary branches. b) The construction process of the evolution network using the neighbor-joining method. The connections were generated between real nodes (genomes or genome patterns) and no bifurcate limitation.

**Figure S2.**
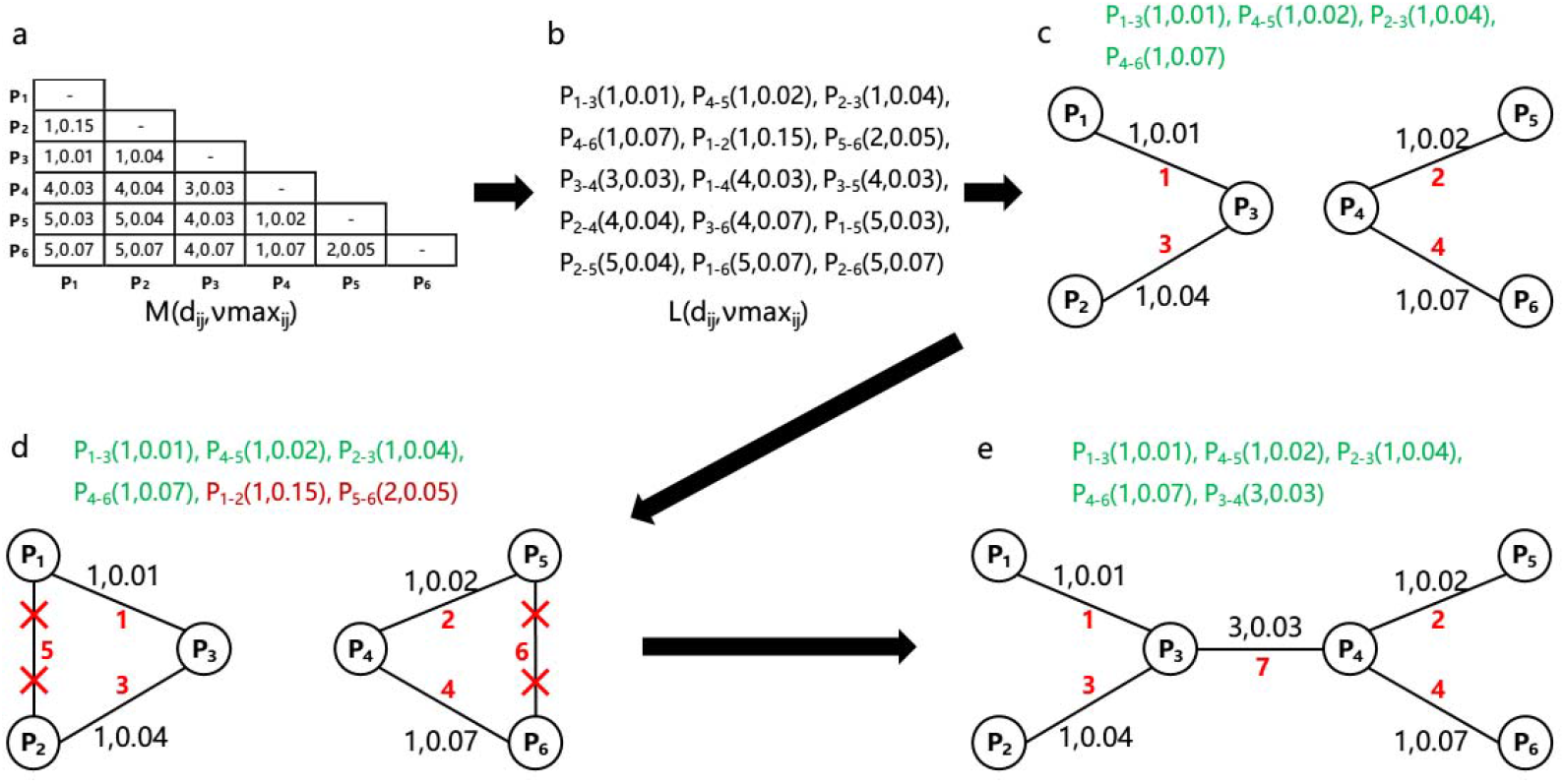
The computing pipeline of VENAS network construction. a) Computing the Hamming distance matrix M(d_ij_, νmax_ij_) adjusted by the max MAF (νmax_ij_) between two genome types; b) Sorting the Hamming distance matrix to an ordered Hamming distance queue L(d_ij_, νmax_ij_); c) Connecting the pairs of genome types in the order of L(d_ij_, νmax_ij_); d) Abandoning the candidate link if there is an indirect connection between the two genome types; e) Abandoning all remaining candidate links in L(d_ij_, νmax_ij_) when the evolution network was fully-connected.

**Figure S3.**
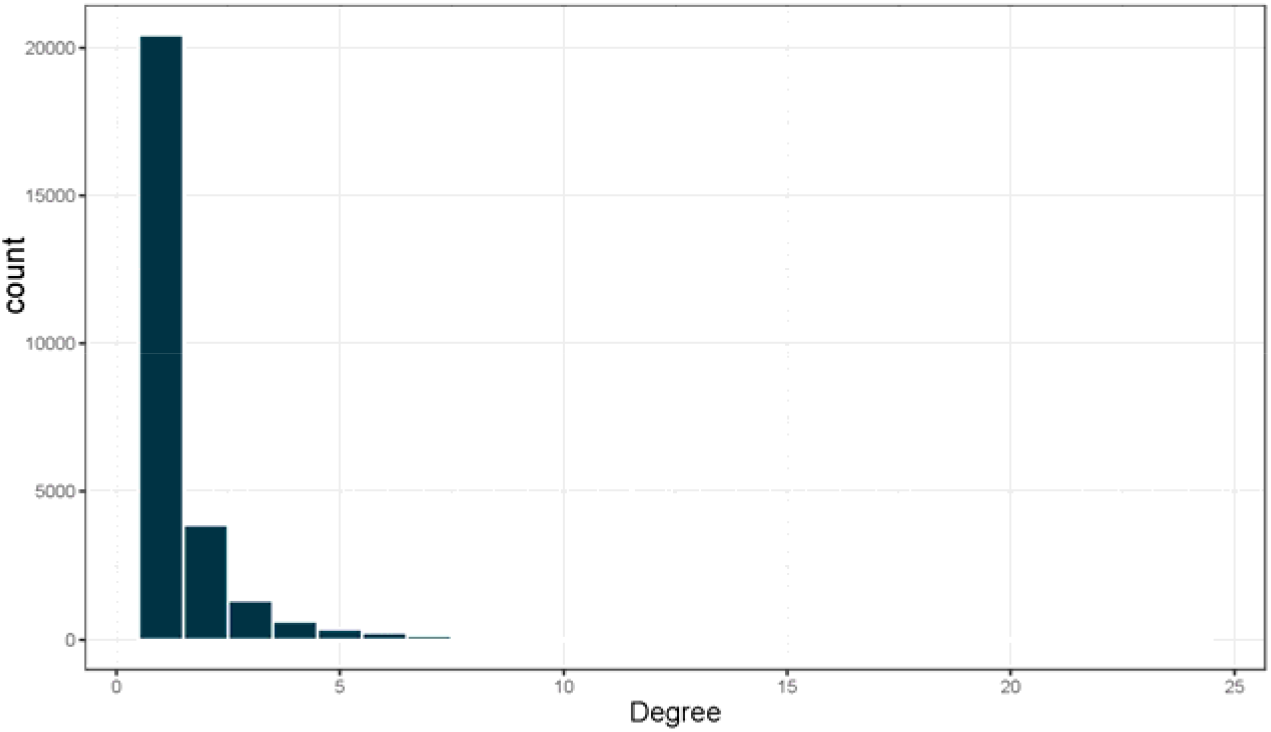
Statistical distribution of the degrees of all nodes in VENAS network.

